# Neural correlates of emotional responses to self-selected music: evidence from multivariate pattern analysis

**DOI:** 10.1101/2025.07.28.667190

**Authors:** Alexandre Sayal, Teresa Sousa, César F. Lima, Miguel Castelo-Branco, Inês Bernardino, Bruno Direito

## Abstract

Music is a uniquely powerful stimulus for evoking complex and deeply felt emotions. While previous research has identified neural correlates of music-evoked emotional responses, less is known about how these felt emotions are represented in the brain, particularly when elicited by familiar, personally meaningful music. Here, we used a personalized fMRI paradigm in which participants (N = 20) each selected musical excerpts corresponding to the nine emotion categories defined by the Geneva Emotional Music Scale. These self-selected excerpts were presented during functional MRI scanning. We first examined the neural correlates of music-evoked emotion by comparing brain activity during music listening to that during exposure to white noise. The maps were consistent with previous research, highlighting clusters in sensory and limbic regions. We then used multivoxel pattern analysis to decode emotion categories from whole-brain activation patterns. The results revealed that music-evoked emotions could be reliably discriminated based on distributed neural activity, with consistent involvement of the superior temporal gyrus, supplementary motor area, amygdala, and cerebellum, among other auditory, motor, and interoceptive regions. These findings provide new insight into the neural encoding of musical emotions and highlight the value of personalized, music-based paradigms for research in auditory and affective neuroscience.

## 1 Introduction

Music elicits a wide range of affective responses, from general feelings of pleasure and arousal to specific emotions like joy, sadness, or nostalgia. The psychological and neural mechanisms underlying these responses are central questions in affective neuroscience (Vuust et al., 2022). Unlike survival-driven emotions, music-evoked emotions are often seen as abstract and deeply personal, offering insight into broader affective processes and potential applications in therapy and neurotechnology (Sayal et al., 2025).

Research on music and emotion distinguishes between perceived emotions - those recognized in music- and felt emotions, which correspond to the subjective experience of the listener (Schubert, 2013). Previous work, including our own, has focused primarily on decoding perceived emotions, identifying neural patterns that correlate with the arousal and valence of musical stimuli (Hoefle et al., 2018; Sayal et al., 2024). In this study, we focus on felt emotions, seeking to decode the neural representations of personal, aesthetically driven affective experiences. This perspective is important, as felt emotions probe the dopaminergic system (Salimpoor et al., 2015) and are highly individualized, influenced by familiarity, personal significance, and prior experience. Familiarity has been shown to enhance emotional engagement with music, leading to stronger activation in the brain’s reward system (C. S. Pereira et al., 2011), thalamus, and superior frontal gyrus (Freitas et al., 2018). By allowing participants to select familiar, personally meaningful musical stimuli, we aimed to maximize the intensity and ecological validity of the experienced emotions (Blood & Zatorre, 2001; Brattico et al., 2011; Tervaniemi, 2023).

Multivoxel pattern analysis (MVPA) has revealed that music-evoked emotions are associated with distributed activity across networks linked to auditory, motor, and interoceptive processing. For example, (Sachs et al., 2018) demonstrated that neural responses to vocal and instrumental emotional sounds expressing happiness, sadness, or fear could be predicted from functional magnetic resonance imaging (fMRI) data, with representations in the primary and secondary auditory cortex, posterior insula, and parietal operculum. Similarly, (Putkinen et al., 2021) found that the auditory and motor cortices reliably discriminated between four music-induced emotions - happiness, sadness, fear, and tenderness - while additional involvement of the somatosensory cortex, insula, and cingulate gyrus suggested a broader network integrating sensory and interoceptive information. (Koelsch et al., 2021) showed that joy and fear were represented in the granular insula, cingulate cortex, premotor and somatosensory cortices, frontal operculum, and auditory cortex, with emotion-related patterns strengthening over the stimulus duration. Complementing these findings, (Trost et al., 2012) identified distinct neural correlates of high- and low-arousal emotions, with high-arousal states engaging sensory and motor regions, and low-arousal states recruiting the ventromedial prefrontal cortex and hippocampus. In sum, these studies show that emotional responses to music can be mapped and decoded using MVPA, although they tend to include a restricted number of emotions.

To structure musical emotions, we adopt the Geneva Emotional Music Scale (GEMS), a tool designed to capture the aesthetic emotions evoked by music, recognizing that musical experiences often elicit emotions distinct from those studied in general affective research (Zentner et al., 2008). This model organizes nine music-specific emotions into three clusters (Figure 1): (1) Sublimity, which includes feelings of wonder, transcendence, tenderness, nostalgia, and peacefulness; (2) Vitality, encompassing emotions of power and joyful activation; and (3) Unease, which includes feelings of sadness and tension. Unlike the emotion models that focus on basic affective dimensions (e.g., valence and arousal), GEMS provides a framework tailored to the complexity of music-induced emotions, offering a tool that is useful to advance perspectives on how these states are represented in the brain.

**Figure 1.**
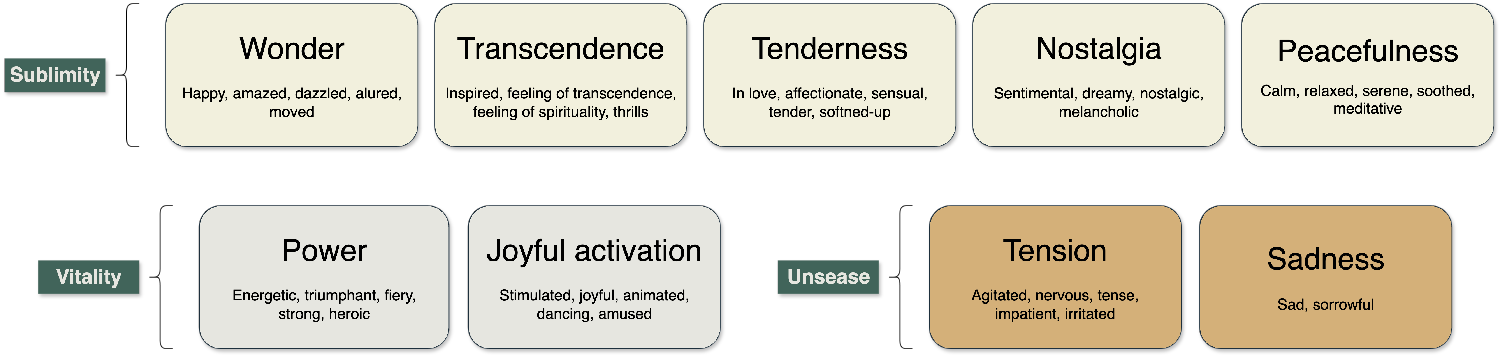
The nine aesthetic emotions framework according to (Zentner et al., 2008).

This study aims to characterize the neural representations of nine aesthetic emotions elicited by self-selected musical stimuli, using fMRI. First, we created a paradigm that, by using self-selected familiar stimuli, maximizes the nine dimensions of the GEMS evoked-emotions model. Second, we map the neural correlates of listening and experiencing these self-selected stimuli. Next, using MVPA, we test whether the nine emotions can be discriminated based on patterns of brain activity. Finally, by analysing the classifier’s parameters, we identify the specific brain regions that most contribute to this discrimination.

Understanding and decoding the neural substrates of music-evoked emotions not only adds to our knowledge of music’s impact on the brain but may also help establish mechanistically valid targets for the development of personalized music-based brain-computer interfaces (Sayal et al., 2025).

## 2 Methods

### 2.1 Participants

Twenty adults (12 females; age range 22-41 years, M = 32.1 years, SD = 6.4 years) participated in the experiment. All participants gave written informed consent. The study was conducted following the Declaration of Helsinki and approved by the Comissão de Ética e Deontologia da Investigação da Faculdade de Psicologia e Ciências de Educação da Universidade de Coimbra. Exclusion criteria were reported diagnoses of neurological, mood, or hearing disorders.

The mood states of the participants were characterized using the Profile of Mood States (POMS) questionnaire (Faro Viana et al., 2001), which includes 42 words describing sensations that people feel in everyday life. Participants were asked to select an answer based on what best corresponds to the way they have been feeling - a scale from 0 (Not at all) to 4 (Very much). Mood state was analyzed according to six subscales - Tension, Depression, Hostility, Vigor, Fatigue, and Confusion - both before and after the acquisition. The average scores for each of the six subscales were 3.34 for Tension, 1.91 for Depression, 1.5 for Hostility, 5.69 for Fatigue, 11.53 for Vigor, 4.75 for Confusion, and there was no difference in the Total Mood Disturbance score from before to after the study (p = 0.11, Supplementary Figure S1).

Our sample reported on average 2.2 ± 3.4 years of formal musical training (range 0-12 years) and scored an average of 18.4 ± 5.1 in the Mini-PROMS (Profile of Music Perception Skills) test (Law & Zentner, 2012; Zentner & Strauss, 2017), a brief, standardized assessment of music perception abilities. It evaluates four key domains: melody discrimination, pitch tuning accuracy, rhythm perception, and tempo detection. The test provides both subscale scores and an overall score, making it a useful tool for quantifying musical aptitude in both musicians and non-musicians.

### 2.2 Stimuli

Participants were asked to select two songs they knew well that evoked each of the nine emotions in the GEMS model. The GEMS was briefly explained, with examples illustrating each emotional category. For instance, the ‘wonder’ emotion factor included terms such as happy, amazed, dazzled, allured, and moved. Participants could select any music, as long as it was available on Spotify.

The instruction was as follows: “Please select, entirely at your discretion, two musical excerpts for each of the nine emotions. Each excerpt should clearly and consistently evoke the intended emotion throughout the 24-second duration. Use Spotify to identify and select the excerpts, indicating the song title and the exact start time in seconds in the table below. Only choose excerpts from music that you already know and/or are familiar with.” A table with the song selection and the timestamps was created for each participant to act as input for the stimulus presentation code. The names and artists of all the selected songs are provided in Supplementary Table T1, showcasing the highly diverse selection across participants.

### 2.3 Experimental Design

#### 2.3.1 Paradigm

Participants listened to their personalized, self-selected musical stimuli, interleaved with white noise. Each trial followed the structure illustrated in Figure 2: 12 seconds of soothing white noise, followed by a 24-second excerpt corresponding to a specific target emotion, a 6-second white noise segment, a second musical 24-second excerpt evoking the same emotion, and finally, 18 seconds of white noise. The target emotion varied across trials, and its order was randomized between runs. In every run, nine unique trials, corresponding to the nine emotion categories, were each presented twice, resulting in 18 trials per run. The scanning session consisted of four runs, each lasting 11 minutes.

**Figure 2.**
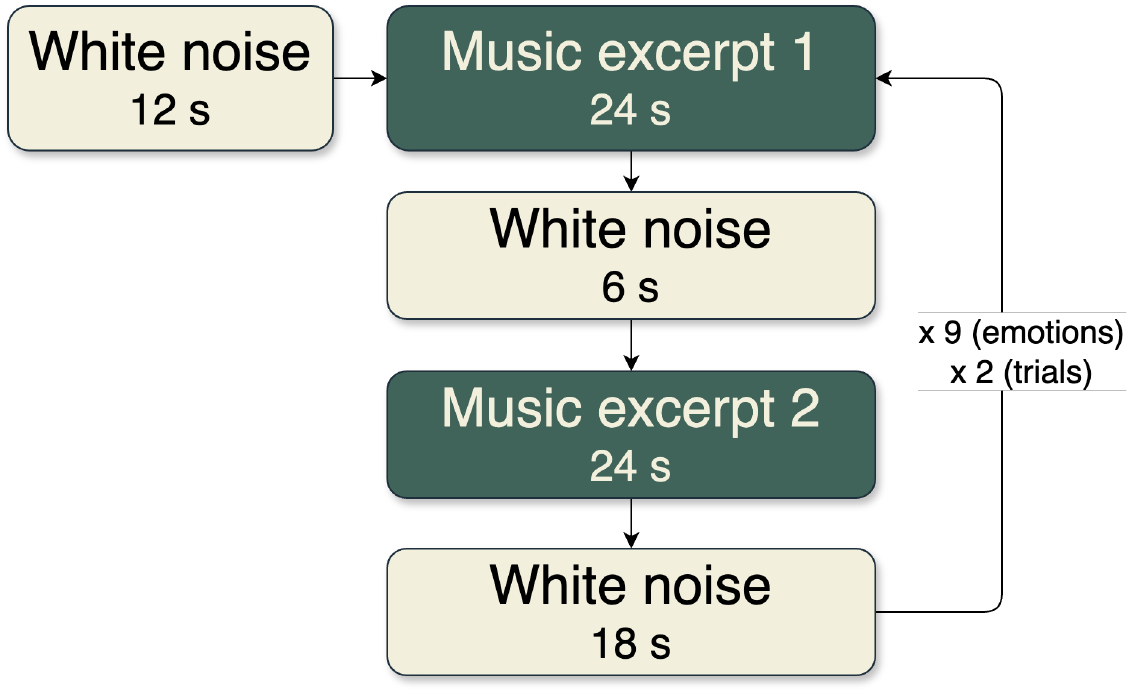
Trial structure of the fMRI paradigm. Each trial was repeated twice for each of the nine emotions.

#### 2.3.2 Implementation

A custom Python-based script was developed to control stimulus presentation. Implemented as a Jupyter Notebook, this script automated the selection and playback of musical stimuli via Spotify’s API (using the package spotipy (Spotipy contributors, 2024)), randomized the emotions per run, and ensured accurate stimulus presentation after receiving the MRI trigger at the beginning of the acquisition.

### 2.4 fMRI Data Acquisition

MRI data were acquired using a 3 T Siemens Magnetom Prisma fit scanner with a 20-channel head coil at the Institute of Nuclear Sciences Applied to Health, Coimbra. Auditory stimuli were presented through MRI-compatible headphones with active noise cancellation (Optoacoustics Optoactive II). The scanning session started with the acquisition of two high-resolution anatomical images: a 3D T1-weighted magnetization-prepared rapid acquisition gradient echo (MPRAGE) sequence (repetition time (TR) = 2530 ms, echo time (TE) = 3.5 ms, flip angle (FA) = 7°, 192 slices, voxel size 1.0 × 1.0 × 1.0 mm, field of view (FOV) = 256 × 256 mm), followed by and a T2-weighted space sequence (TR = 3200 ms, TE = 408 ms, 192 slices, voxel size 1.0 × 1.0 × 1.0 mm, FOV = 250 × 250 mm). Subsequently, four functional runs were acquired using a 2D multi-band (MB) gradient-echo echo-planar imaging (EPI) sequence (TR = 1000 ms, TE = 37 ms, flip angle = 68°, 600 volumes, 66 slices, voxel size 2.0 × 2.0 × 2.0 mm, FOV = 220 mm, MB factor = 6). To correct for image distortions caused by magnetic field inhomogeneities, we acquired pairs of spin-echo images with anterior-posterior and posterior-anterior phase encoding polarity with matching geometry and echo-spacing to each of the functional scans (TR = 10250 ms, TE = 73.6 ms). These images were acquired before the functional runs.

### 2.5 fMRI Data Analysis

The MRI data were organized according to the Brain Imaging Data Standard (BIDS), using a customizable conversion tool (Tyszka, 2023) for the DICOM images and custom scripts to associate the task events. Preprocessing was performed in fMRIPrep v23.1.2 (Esteban et al., 2019, 2023) and included slice time correction, motion correction, unwarping, and normalization to the MNI space. See Supplementary Materials for a complete description of the fMRIPrep methods.

The following sections describe the imaging data analyses performed using custom Python scripts based on the package Nilearn v0.11.1 (Abraham et al., 2014; Nilearn contributors et al., 2025).

#### 2.5.1 GLM Analysis

First-level analyses were performed using a single General Linear Model (GLM) estimated separately for each participant. The design matrix included one predictor for each of the nine emotion categories, based on the participant’s self-labeled musical stimuli, as well as nuisance regressors to account for head motion (six motion parameters, their first-order derivatives, and squared terms, totaling 24 confound regressors). Prior to model estimation, we applied a high-pass temporal filter with a cut-off frequency of 0.007 Hz and spatial smoothing using a Gaussian kernel with a full-width at half-maximum (FWHM) of 4 mm. A second-order autoregressive model (AR(2)) was used to account for temporal autocorrelations.

For the second-level analysis, we computed a one-way ANOVA with ‘emotion’ as the within-subjects factor. This analysis used the first-level contrast maps for each emotion (emotion *>* white noise) to identify brain regions where the magnitude of responses varied as a function of emotion. We restricted our analysis to grey-matter voxels, defined using the MNI152NLin2009cAsym segmentation probability map with a threshold of 0.15. To correct for multiple comparisons, we applied Bonferroni correction at the voxel level (p *<* 0.05). Clusters were defined using a minimum cluster size of 25 contiguous voxels. Anatomical labels for significant clusters were assigned based on the AAL3 maps (Rolls et al., 2020) and cross-validated with functional annotations from the Neurosynth database (Yarkoni et al., 2011).

#### 2.5.2 Multivoxel Pattern Analysis

We performed feature selection to focus the analysis on a subset of voxels most likely to provide discriminative information. Specifically, we used a voxel-wise stability metric as described by (Just et al., 2010). This approach identifies voxels whose activation profiles across the nine emotions were consistent over repeated presentations of the musical excerpts, i.e., indicating stable relative variation between emotional responses. The rationale is that stable voxels are more likely to reflect meaningful, stimulus-driven neural responses. The voxel stability was estimated as the average pairwise Pearson correlation between its nine-emotion activation profiles, a vector of nine responses of a voxel to the emotions during a single presentation, across repeated presentations of the nine emotion categories. Voxels with an average stability correlation of at least 0.1 were retained, resulting in individual stability masks for each participant. This threshold, while defining a large feature input space, was chosen as a compromise to provide a clear criterion for stability while ensuring a comprehensive sampling of the whole brain and imposing minimal restrictions.

For each 24-second music excerpt, we extracted denoised BOLD signals after regressing out nuisance variables related to head motion (six motion parameters, their first-order derivatives, and squared terms, totaling 24 confound regressors), applying a high-pass temporal filter with a cut-off frequency of 0.007 Hz, detrending, and normalizing the time series to unit variance. Each excerpt was divided into four consecutive, non-overlapping 6-second segments. The BOLD signal within each segment was averaged, resulting in four time-resolved feature values per voxel per block. Feature extraction was restricted to gray matter voxels, defined using the MNI152NLin2009cAsym gray matter probability map thresholded at 0.15.

Multivariate pattern analysis was conducted using a supervised learning method as the estimator model - a Support Vector Machine (SVM) with a linear kernel and an L1 regularization factor. The multiclass problem was addressed using a one vs. all method. We implemented cross-validation to split the data into different sets. We then fitted the estimator on the training dataset and measured an unbiased error on the testing set. To avoid temporal dependencies between training and testing samples (bias due to temporal proximity of samples, particularly relevant in functional data), we implemented a leave-one-run-out (LORO) strategy, where the classifier was trained on three of the four runs and tested on the remaining one. Leaving out blocks of correlated observations, rather than individual observations, is crucial for non-biased estimates (Varoquaux et al., 2017).

## 3 Results

Our results are presented in two parts. We first mapped the brain responses to music across emotions (univariate analysis), and then assessed the discriminability of music-evoked emotions using MVPA and explored the spatial distribution of the most discriminative voxels across the brain.

### 3.1 Brain network of music-evoked emotions

Figure 3 displays brain regions where activity varied significantly as a function of emotion (i.e., main effect of emotion), based on whole-brain analysis. Activations were largely bilateral, including highly significant clusters in the superior temporal gyrus, Heschl’s gyrus, the cerebellum, supplementary motor area, putamen, insula, frontal cortex, and nucleus accumbens (Table 1).

**Table 1:** Cluster table of the activation map of Figure 3. MNI Coordinates of the peak voxel, the z-value of the peak voxel, the cluster size, and the label from the AAL3 atlas.

| Cluster label | X | Y | Z | Z-Value of Peak Voxel | Cluster Size ($mm^3$) |
| --- | --- | --- | --- | --- | --- |
| Temporal_Sup_R | 54 | 2 | -5 | 18.45 | 22224 |
| Temporal_Sup_L | -53 | -15 | 6 | 17.82 | 22352 |
| Cerebellum_6_R | 30 | -61 | -25 | 14.41 | 8104 |
| Cerebellum_6_L | -33 | -63 | -23 | 13.27 | 2728 |
| Precentral_R | 56 | -3 | 46 | 12.05 | 2160 |
| Supp_Motor_Area_L | -1 | 2 | 68 | 11.49 | 3752 |
| Postcentral_L | -55 | -7 | 48 | 11.37 | 1864 |
| Parietal_Inf_R | 60 | -53 | 40 | 10.71 | 10408 |
| Cingulate_Mid_R | 4 | -33 | 48 | 10.69 | 22528 |
| Frontal_Sup_2_R | 28 | 34 | 50 | 10.37 | 15008 |
| Cerebellum_8_L | -25 | -65 | -53 | 9.76 | 1568 |
| ACC_pre_R | 4 | 46 | 12 | 9.21 | 6880 |
| Supp_Motor_Area_L | -9 | 14 | 70 | 9.15 | 928 |
| Temporal_Mid_R | 68 | -23 | -13 | 8.80 | 1960 |
| Cerebellum_Crus1_L | -49 | -65 | -27 | 8.55 | 344 |
| Cerebellum_8_R | 18 | -69 | -53 | 8.54 | 2712 |
| Cerebellum_Crus2_L | -49 | -61 | -45 | 8.51 | 1712 |
| Cerebellum_7b_L | -15 | -79 | -45 | 8.21 | 552 |
| Insula_R | 34 | 18 | -11 | 8.16 | 440 |
| Occipital_Sup_L | -19 | -67 | 26 | 8.08 | 1152 |
| Temporal_Inf_R | 48 | 10 | -41 | 8.02 | 1240 |
| Cerebellum_6_L | -11 | -67 | -13 | 7.89 | 824 |
| Putamen_R | 24 | 4 | 6 | 7.84 | 544 |
| Temporal_Inf_R | 58 | -7 | -35 | 7.72 | 1072 |
| Angular_R | 46 | -77 | 40 | 7.65 | 3456 |
| Temporal_Inf_L | -53 | -9 | -31 | 7.65 | 992 |
| Insula_R | 40 | -11 | -1 | 7.45 | 216 |
| Frontal_Inf_Tri_R | 54 | 36 | 4 | 7.24 | 632 |
| N_Acc_R | 14 | 16 | -11 | 7.18 | 272 |
| Supp_Motor_Area_R | 14 | 20 | 62 | 7.02 | 400 |
| Broca_R | 30 | 30 | 4 | 6.85 | 216 |
| Occipital_Mid_L | -37 | -83 | 38 | 6.77 | 296 |
| Postcentral_R | 54 | -7 | 26 | 6.77 | 408 |
| Frontal_Mid_2_L | -27 | 18 | 60 | 6.75 | 208 |
| Frontal_Mid_2_L | -47 | 48 | -5 | 6.51 | 240 |
| N_Acc_L | -11 | 18 | -9 | 6.23 | 224 |
| Frontal_Sup_2_R | 26 | 64 | -3 | 6.08 | 288 |
| Temporal_Pole_Mid_L | -53 | 10 | -29 | 5.91 | 256 |
| Angular_L | -61 | -59 | 22 | 5.81 | 216 |

**Figure 3.**
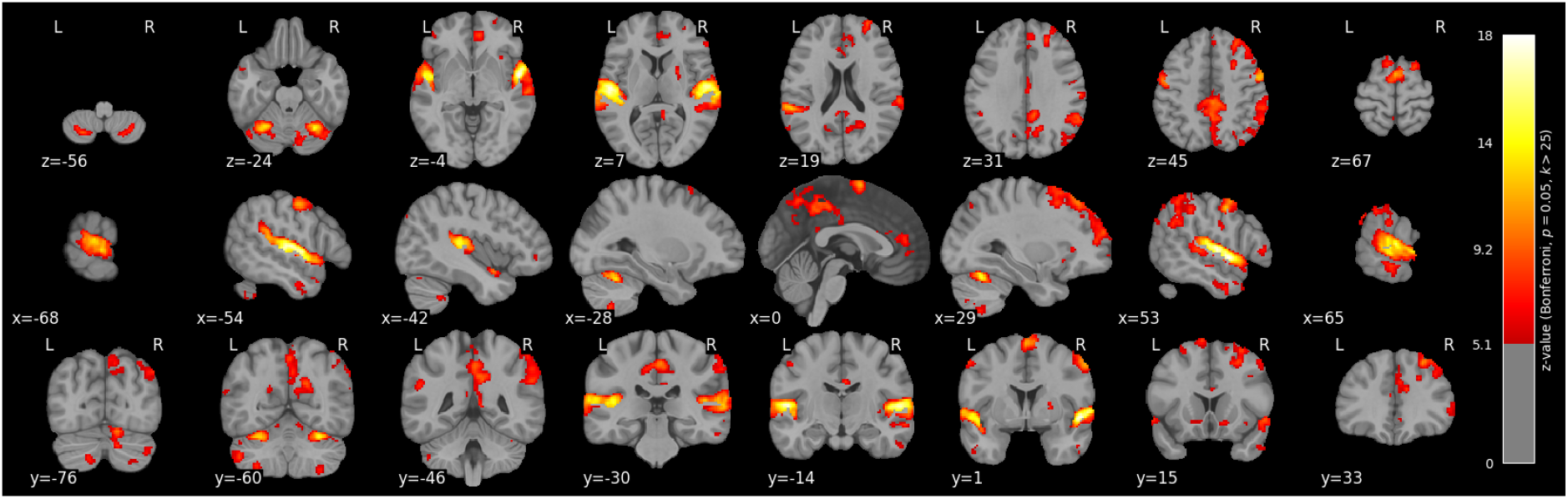
Brain regions in which BOLD responses varied as a function of music-evoked emotions. The map shows z-values corrected for multiple comparisons with Bonferroni’s method (p = 0.05) and a minimum cluster size of 25 voxels.

### 3.2 Decoding music-evoked emotions

To test whether the nine aesthetic emotions could be reliably discriminated based on neural activation patterns, we applied an MVPA pipeline to whole-brain fMRI data. Feature selection was performed using a stability-based mask, ensuring that only voxels with consistent emotion-related activation profiles contributed to the analysis. The linear SVM classifier was trained and tested using a leave-one-run-out cross-validation scheme. Classification accuracy served as the performance metric, with a chance level of 11.1% given the nine emotion classes. This analysis assessed whether distributed patterns of brain activity contained sufficient information to predict the specific emotion class evoked by music.

#### 3.2.1 Voxel selection

In Figure 4, we display the sum of the stability masks estimated for each subject. Clusters were found in the temporal superior gyrus, precentral and postcentral gyrus, supramarginal gyrus, and the SMA. The average number of voxels in the stability mask across subjects was 8205.6 ± 2129.6 voxels.

**Figure 4.**
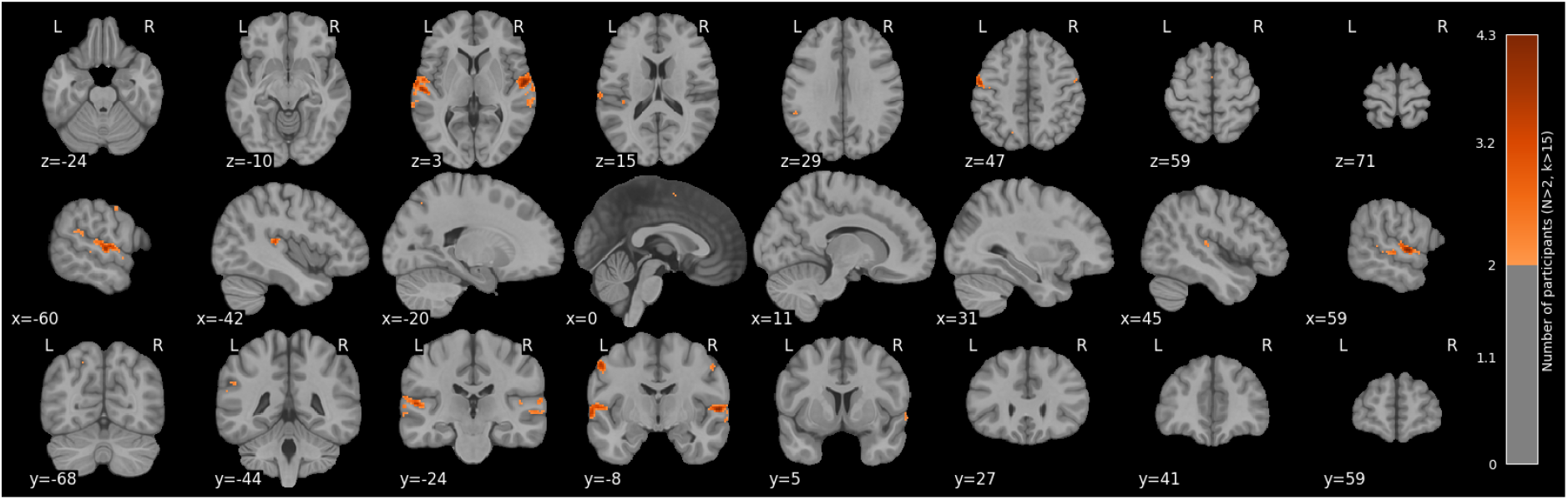
Sum of the binary stability masks for the 20 subjects. For this map, we set the threshold to a minimum of 2 participants, with cluster correction of k*>*5 voxels and spatial smoothing (FWHM = 4 mm).

#### 3.2.2 Classification performance

Overall, our model achieved 48.5% ± 6.2% accuracy across all emotions. In Figure 5, we display the confusion matrix with the average percentages attributed to each class, where a clear discrimination between the correct classification (diagonal) is visible, with no major heterogeneity in the incorrect classifications.

**Figure 5.**
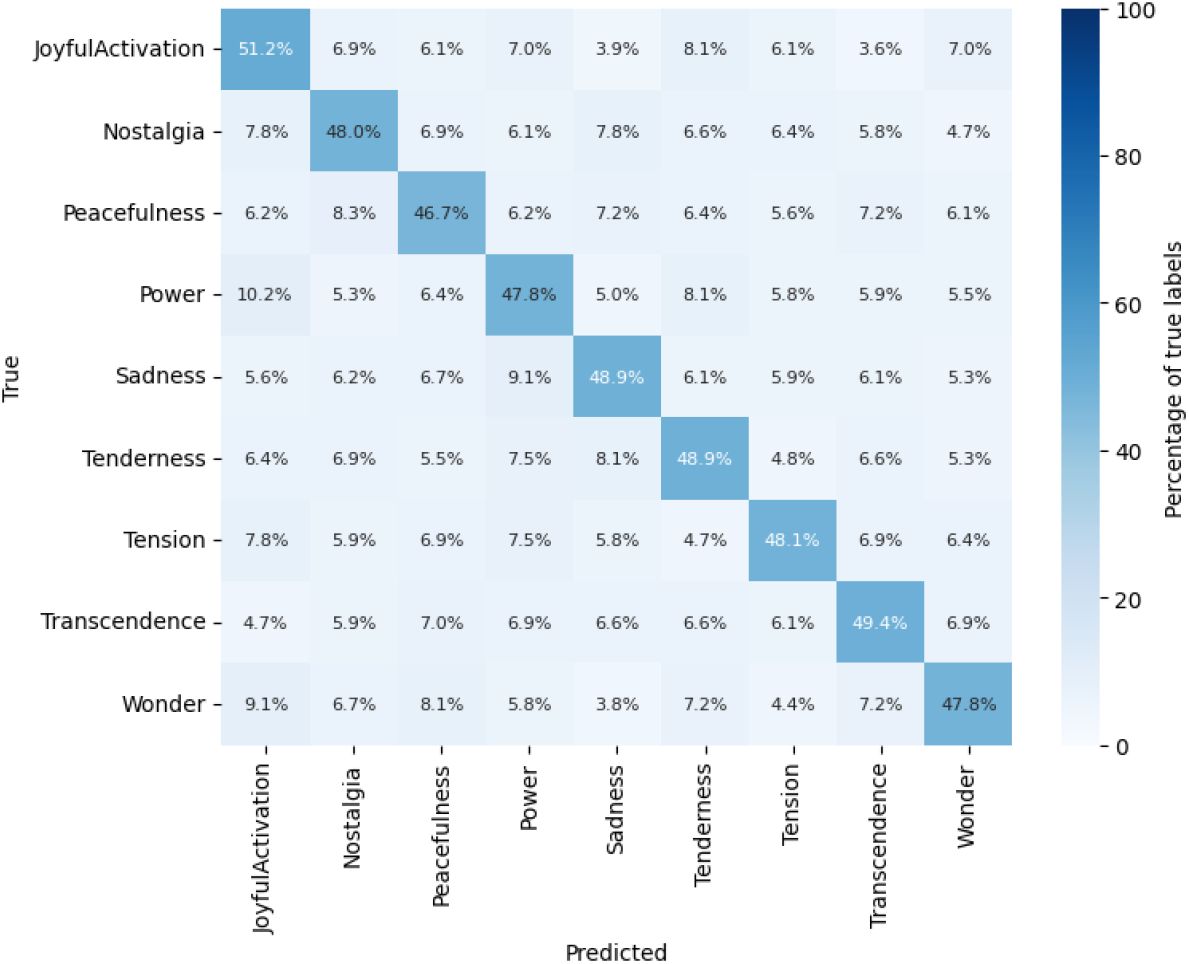
Group-level confusion matrix showing the results of the testing phase across the 20 participants. The overall balanced accuracy was 48.5 ± 6.2%, while the chance level sits at 11.1% in this case.

#### 3.2.3 Discriminative brain regions

From the trained models, we retrieved the classifier’s weights for each voxel and emotion class for each participant. These weights indicate the relative contribution of each voxel to distinguishing a given emotion from the others. To obtain a group-level summary of voxel importance, we summed the absolute weights across all emotions and participants, achieving a predictive power map, displayed in Figure 6. The clusters identified in this map are listed in Table 2.

**Table 2:**
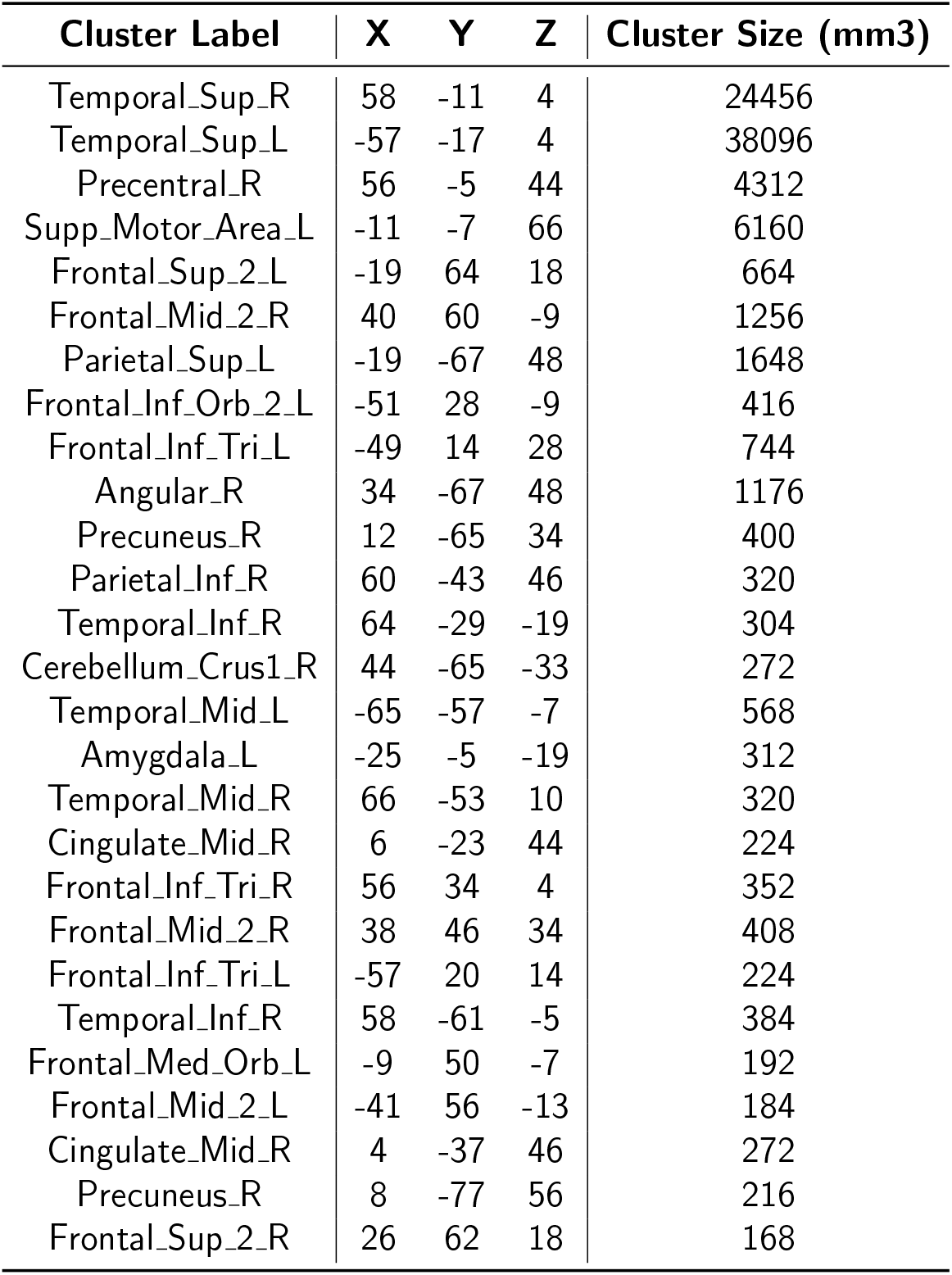
Cluster table of the predictive power map of Figure 6, showing the MNI coordinates of the peaks, cluster size, and AAL3 labeling.

| Cluster Label | X | Y | Z | Cluster Size (mm3) |
| --- | --- | --- | --- | --- |
| Temporal_Sup_R | 58 | -11 | 4 | 24456 |
| Temporal_Sup_L | -57 | -17 | 4 | 38096 |
| Precentral_R | 56 | -5 | 44 | 4312 |
| Supp_Motor_Area_L | -11 | -7 | 66 | 6160 |
| Frontal_Sup_2_L | -19 | 64 | 18 | 664 |
| Frontal_Mid_2_R | 40 | 60 | -9 | 1256 |
| Parietal_Sup_L | -19 | -67 | 48 | 1648 |
| Frontal_Inf_Orb_2_L | -51 | 28 | -9 | 416 |
| Frontal_Inf_Tri_L | -49 | 14 | 28 | 744 |
| Angular_R | 34 | -67 | 48 | 1176 |
| Precuneus_R | 12 | -65 | 34 | 400 |
| Parietal_Inf_R | 60 | -43 | 46 | 320 |
| Temporal_Inf_R | 64 | -29 | -19 | 304 |
| Cerebellum_Crus1_R | 44 | -65 | -33 | 272 |
| Temporal_Mid_L | -65 | -57 | -7 | 568 |
| Amygdala_L | -25 | -5 | -19 | 312 |
| Temporal_Mid_R | 66 | -53 | 10 | 320 |
| Cingulate_Mid_R | 6 | -23 | 44 | 224 |
| Frontal_Inf_Tri_R | 56 | 34 | 4 | 352 |
| Frontal_Mid_2_R | 38 | 46 | 34 | 408 |
| Frontal_Inf_Tri_L | -57 | 20 | 14 | 224 |
| Temporal_Inf_R | 58 | -61 | -5 | 384 |
| Frontal_Med_Orb_L | -9 | 50 | -7 | 192 |
| Frontal_Mid_2_L | -41 | 56 | -13 | 184 |
| Cingulate_Mid_R | 4 | -37 | 46 | 272 |
| Precuneus_R | 8 | -77 | 56 | 216 |
| Frontal_Sup_2_R | 26 | 62 | 18 | 168 |

**Figure 6.**
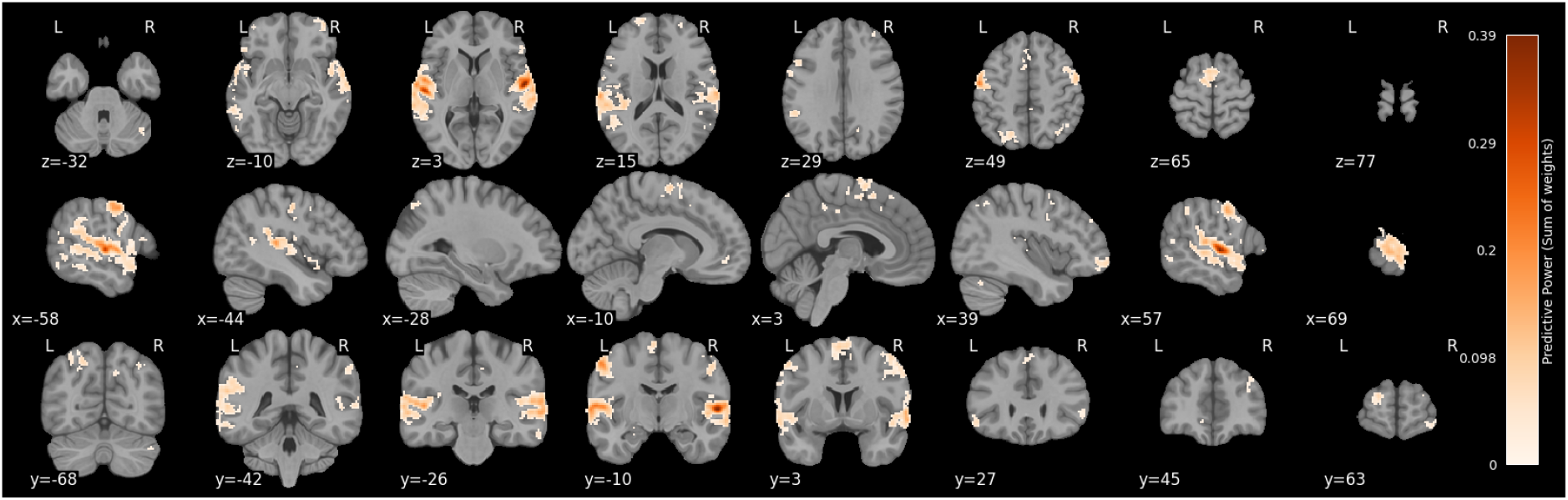
Predictive power brain map for all conditions and subjects. The map shows the sum of the absolute weights of the linear SVM classifier across all participants and emotions. The color bar indicates the relative contribution of each voxel to emotion classification.

## 4 Discussion

This study investigated the neural representation of music-evoked emotions using a personalized fMRI paradigm and multivariate decoding. Participants selected familiar musical excerpts evoking the nine emotion factors of the Geneva Emotional Music Scale, allowing us to maximize emotional salience and ecological validity. Univariate analyses revealed emotion-specific neural activity across a broad bilateral network. Multivariate analyses using MVPA then showed that these emotions could be reliably decoded from whole-brain activity using a linear SVM classifier, with performance significantly above chance. Brain regions contributing to classification included auditory, motor, and interoceptive areas, consistent with prior studies linking these systems to music-induced affective processing (Koelsch, 2020; Vuust et al., 2022).

The successful decoding of emotional categories reinforces findings from previous MVPA studies that mostly explored different emotions across the valence dimension (e.g., joy vs. sadness) (Koelsch et al., 2021; Putkinen et al., 2021) and supports the view that aesthetic emotions are represented in distributed networks. Here, we explore the full spectrum of the GEMS, which we considered an important advancement when studying the correlates of felt emotions. The use of self-selected, familiar music departs from more controlled but less engaging experimental approaches and shifts the focus from perceived to felt emotions. Familiarity has been shown to enhance emotional engagement and activity in reward-related areas (Freitas et al., 2018; C. S. Pereira et al., 2011), and our paradigm leveraged this to enhance the ecological validity and intensity of the emotional experience.

Our univariate analysis replicated earlier work (Koelsch, 2020; Trost et al., 2012), showing broad activation in sensory and limbic regions during music listening compared to noise. These results suggest that participant-specific emotional experiences, despite being highly individualized, are underpinned by consistent neural mechanisms. Importantly, we also explored emotion-specific activation patterns, but found that most of the core clusters, especially in auditory, motor, and reward-related regions, responded similarly across all emotional categories (Supplementary Figure S4). This lack of fine-grained selectivity supports the idea that these areas act not as “emotion-specific modules” but as part of a general affective interpretation network, consistent with the theory of constructed emotion (Barrett, 2016, 2017), which posits that emotions emerge from domain-general systems involved in conceptualization and interoception.

Despite the lack of strong univariate selectivity, we were still able to successfully discriminate emotion categories using multivoxel pattern analysis. This reinforces the view that emotional states are encoded not through isolated regional activations but through distributed spatial patterns across multiple brain systems. This observation also aligns with emerging evidence from connectivity-based studies, which emphasize the importance of network-level integration in affective processing (Kober et al., 2008; Lindquist et al., 2016; Pessoa, 2017). Emotion regulation and differentiation appear to depend not only on activity within specific nodes but on the functional coupling between regions involved in perception, interoception, memory, and conceptualization. Gaining finer control over this network-level architecture may be essential for both mechanistic understanding and music-based emotion regulation interventions.

Our task design demonstrates that it is possible to obtain reliable univariate and multivariate results when using individualized, self-selected music and focusing specifically on felt emotional experiences. This personalized approach not only enhances the ecological validity of studies on musical emotions, a recognized need in music research (Tervaniemi, 2023), but also holds promise for applied fields such as music-based rehabilitation and brain-computer interfaces. These interventions could be optimized for effectiveness by tailoring musical stimuli to individual preferences and emotional responses.

### 4.1 Limitations and Future Directions

While our personalized paradigm enhances emotional engagement and ecological validity, it also introduces methodological challenges. Among these is the large between-subject heterogeneity in the selected musical stimuli. Participants’ choices varied widely in terms of low-level acoustic features (e.g., tempo, timbre, loudness) and genres. This variability complicates interpretation: although the classifier reliably distinguished between emotion categories, it remains unclear to what extent decoding reflects affective processes as opposed to differences in stimulus properties. It is important to note, however, that the auditory cortex does not merely process sensory/acoustic information, but is functionally embedded within broader emotion-related networks. As highlighted by (Koelsch et al., 2021), the auditory cortex has direct anatomical and functional connections to limbic and paralimbic structures such as the amygdala, insula, cingulate cortex, and ventral striatum, and plays a central role in generating feeling representations from sound. Furthermore, its functional connectivity with reward-related regions has been shown to predict the subjective emotional value of music and to vary with individual traits such as musical anhedonia (Martínez-Molina et al., 2016). Thus, even though our paradigm did not fully control for acoustic features across emotion categories, it is likely that decoding success in auditory areas reflects, at least in part, their involvement in affective appraisal and emotional experience, rather than simply encoding acoustic differences.

The self-selection of stimuli also limits our ability to pinpoint which mechanisms of emotion induction were engaged. According to the BRECVMA model (Juslin, 2013), music can evoke emotions through multiple mechanisms, including brainstem reflexes, rhythmic entrainment, evaluative conditioning, musical expectancy, autobiographical memory, and others. Our personalized paradigm likely triggered a combination of these pathways, particularly those linked to personal associations and memories, yet their individual contributions could not be isolated. Disentangling these pathways will require paradigms specifically designed to manipulate each mechanism independently, possibly through controlled stimulus design or complementary behavioral measures.

Finally, the use of a linear support vector machine as our decoding model represents a methodological trade-off. While linear classifiers offer strong interpretability and are robust to overfitting in highdimensional fMRI data, they may not capture more complex, non-linear relationships between neural activity and emotional states (Hebart & Baker, 2018; F. Pereira et al., 2009). That said, if the feature space is well-structured and informative, as was ensured here through stability-based selection, then classifier complexity may be less critical. Future work could explore the use of non-linear models or deep learning approaches, especially in larger datasets, to determine whether more flexible classifiers can uncover additional structure in the representation of music-evoked emotions.

Looking forward, combining personalized paradigms with computational models of acoustic and emotional features may help reconcile ecological validity with experimental control. Additionally, incorporating realtime decoding techniques could pave the way for music-based neurofeedback or affective brain-computer interfaces, with promising applications in clinical contexts where improving emotion regulation is a key therapeutic target.

## Data and Code Availability

All scripts used for the analyses mentioned in this work can be found in this git repository: https://github.com/CIBIT-UC/brainplaybacktask02. The anonymized and defaced dataset can be found in BIDS format at https://doi.org/10.57979/P4SAYV. The data management plan regarding this project can be found at Zenodo https://doi.org/10.5281/zenodo.10563831.

## Author Contributions

Conceptualization - AS, IB, BD; Data curation - AS, BD, Formal Analysis - AS, BD, Funding acquisition-MCB, IB, BD; Investigation - AS, BD, Methodology - AS, CL, BD; Project administration - IB, BD; Resources - BD, MCB; Software - AS, BD; Supervision - TS, MCB, BD, Visualization - AS, BD, Writing-original draft - AS, BD; Writing - review & editing - AS, CL, IB, TS, MCB, BD

## Acknowledgements

We thank all the participants who took part in this study. We also thank the remaining team members of the Brainplayback project - André Granjo, Carolina Travassos, João Pereira, Renato Panda, Rui Pedro Paiva, Daniela Pereira, and Inês Almeida - for the insightful contributions. A last word of gratitude to the team of the MRI Unit of ICNAS - Sónia Afonso, Tânia Lopes - for their support and assistance during data acquisition.

