## Supplementary Materials for "Neural correlates of emotional responses to self-selected music: evidence from multivariate pattern analysis"

#### POMS scores before vs. after the MRI session

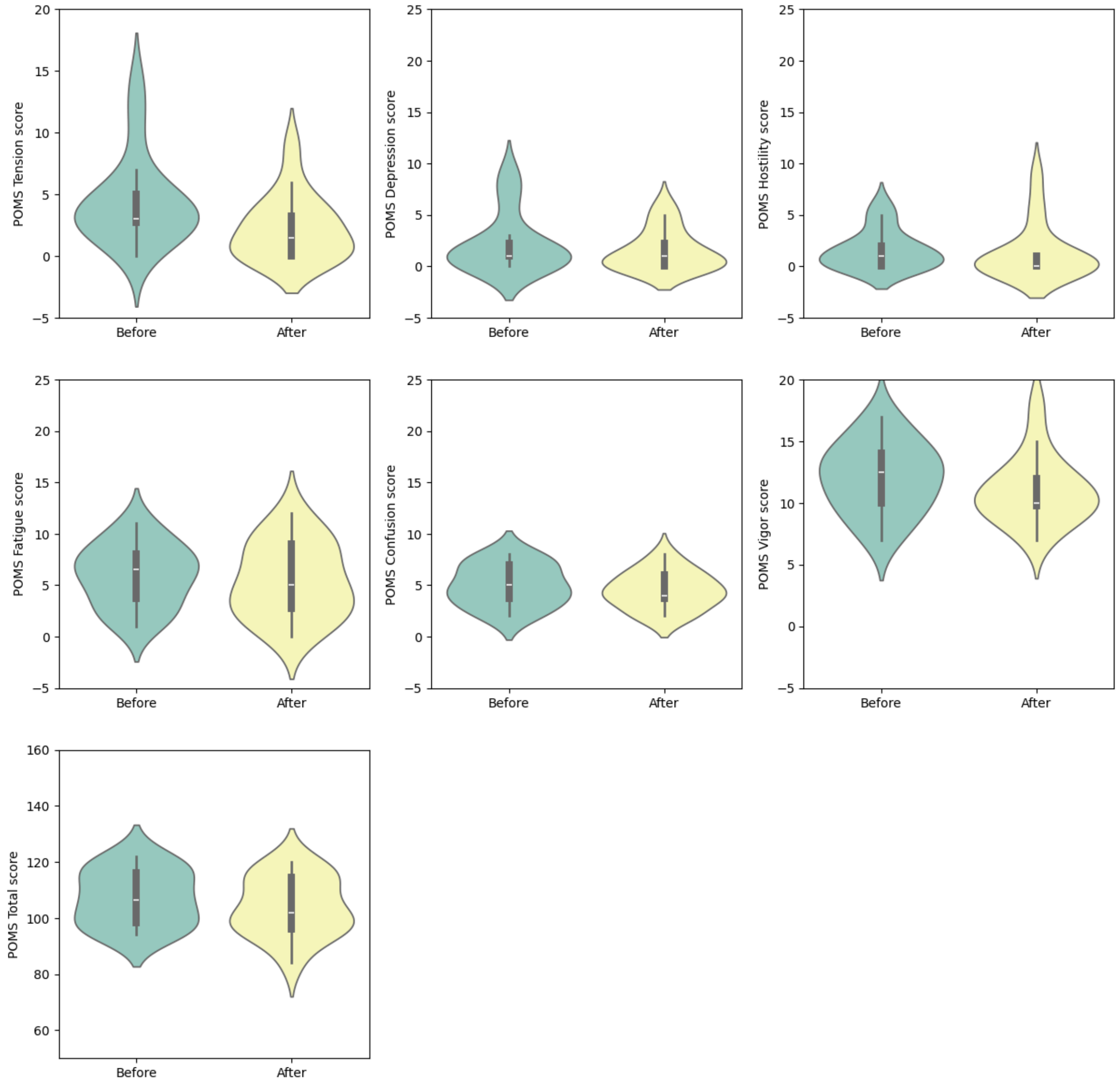


**Figure S1 - Average POMS scores across participants before and after the MRI session. Using Wilcoxon's paired sample test with Bonferroni's correction, no significant differences were found.**

### Self-selected song list per emotion

**Table S1 - Full set of song names and artists selected by the participants, organized per emotion.**

| Emotion | Song Name | Artist |
|---|---|---|
| Joyful activation | Don't Stop Me Now - Remastered 2011 | Queen |
| | Steal My Kisses | Ben Harper And The Innocent Criminals |
| | Never Gonna Give You Up | Rick Astley |
| | Gisela | Bárbara Tinoco |
| | Virtual Insanity | Various Artists |
| | Chicago | Sufjan Stevens |
| | Deixa pra Lá | Bossacucanova |
| | All I Ever Wanted | Hard Lights |
| | Beijo de Funana | Némanus |
| | Tô Voltando | Simone |
| | Proud Mary | Tina Turner |
| | Burra | Karetus |
| | Monsters, Inc. | Randy Newman |
| | Wake Me Up Before You Go-Go | Wham! |
| | I'm Gonna Be (500 Miles) | The Proclaimers |
| | Anda estragar-me os planos | Salvador Sobral |
| | One More Time - Otra Vez | SUPER JUNIOR |
| | Shut Up and Dance | WALK THE MOON |
| | Da Coconut Nut - The Coconut Song | The San Miguel Master Chorale |
| | DANCE DANCE | DAY6 |
| | All My Friends | LCD Soundsystem |
| | Dancing With Myself | Billy Idol |
| | Dancing in the Moonlight | Toploader |
| | Mambeado | Onda Vaga |
| | Hey Ya! | Outkast |
| | Tank! | SEATBELTS |
| | stupid horse | 100 gecs |
| | Bonito | Jarabe De Palo |
| | She's so High | Tal Bachman |
| | I AM | IVE |
| | Another Day Of Sun | Justin Hurwitz |

| | | |
|---|---|---|
| | SAVE The World | Toby Fox |
| | Feel It Still | Portugal. The Man |
| | Imagine | 브런치 |
| | Balalaikas | Anaquim |
| | What You Know | Two Door Cinema Club |
| | Human | The Killers |
| | La Vida Tombola | Manu Chao |
| | Use It | The New Pornographers |
| Nostalgia | Swan Lake Suite, Op. 20a: I. Scene "Swan Theme". Moderato | Pyotr Ilyich Tchaikovsky |
| | Zorro | António Zambujo |
| | Kidult | SEVENTEEN |
| | Rocket Man (I Think It's Going To Be A Long, Long Time) | Elton John |
| | Eram os Teus Olhos | Martim Vicente |
| | Everything In Its Right Place | Radiohead |
| | Dance of the Sugar Plum Fairy | Lior Rosner |
| | By This River - 2004 Digital Remaster | Brian Eno |
| | Sunday Candy | Nico Segal |
| | Tanto Mar | Chico Buarque |
| | Cold | Jorge Méndez |
| | Cinema Paradiso (Main Theme) | Ennio Morricone |
| | Saturn | Sleeping At Last |
| | Menino | Quinta Do Bill |
| | Quero é viver | Humanos |
| | Heat Waves | Glass Animals |
| | Nothing Matters | The Last Dinner Party |
| | Come Here | Kath Bloom |
| | Song for Zula | Phosphorescent |
| | Colors | Black Pumas |
| | Fireflies | Owl City |
| | Golden Brown | The Stranglers |
| | Suite No. 3, Op. 19, No. 1 (version for 2 violins and string orchestra): I. Prelude: Adagio | Henning Kraggerud |
| | Vessel | Jon Hopkins |
| | Extreme Battle The Three Super Saiyans | Dragon Ball |
| | Menino do Bairro Negro | José Afonso |
| | Ele e Ela | Ala Dos Namorados |

| | | |
|---|---|---|
| | Glimpse of Us | Joji |
| | Carga de ombro | Samuel Úria |
| | Somewhere Over The Rainbow | Leanne & Naara |
| | Speed of Sound | Coldplay |
| | See You Again (feat. Charlie Puth) | Wiz Khalifa |
| | Lepo Lepo | Psirico |
| | In A Ocean Of Joy | Dreamcatcher |
| | Ilha de Santiago | Mayra Andrade |
| | If I Fell - Acoustic / Live At The Hit Factory, NYC / 2003 | Maroon 5 |
| | Better Together | Jack Johnson |
| | Sunrise | Norah Jones |
| | Piano Concerto No. 23 in A Major, K. 488: I. Allegro (Cadenza by Busoni) | Wolfgang Amadeus Mozart |
| | Welcome to the Black Parade | My Chemical Romance |
| Peacefulness | Salento | René Aubry |
| | Trafic d'orgasmes | Le Sextet à Claques |
| | Clair de Lune | Johann Debussy |
| | Minefields | Faouzia |
| | (Sittin' On) the Dock of the Bay | Otis Redding |
| | She's Only Happy In The Sun | Ben Harper |
| | Keep Your Head Up | Ben Howard |
| | Gabriel's Oboe | Ennio Morricone |
| | July | Noah Cyrus |
| | The Mummers' Dance | Loreena McKennitt |
| | Servants of the Mountain | SQUARE ENIX MUSIC |
| | In The Waiting Line | Zero 7 |
| | River Flows In You | Yiruma |
| | Breathe | Two Steps from Hell |
| | Climb Together | Audiomachine |
| | Opia | Hedflux |
| | Suite bergamasque, L. 75: III. Clair de lune | Claude Debussy |
| | Heaven Only Knows | Mellah |
| | Space Song | Beach House |
| | Live Your Life | Yuna |
| | Gentle Moon | Sun Kil Moon |
| | Tiger Mountain Peasant Song | Fleet Foxes |

| | | |
|---|---|---|
| | Slow Dancing in a Burning Room | John Mayer |
| | Mercury | Bryce Dessner |
| | Eternal Beauty | Nils Landgren |
| | Lien-yun st. | Isato Nakagawa |
| | ocean eyes | Billie Eilish |
| | Still Broke - A COLORS ENCORE | Samm Henshaw |
| | Love Theme | Ennio Morricone |
| | Flashlight - From "Pitch Perfect 2" Soundtrack | Jessie J |
| | Luz Do Sol | Caetano Veloso |
| | Whitacre: The Seal Lullaby | Eric Whitacre |
| | Shimbalaiê - Live | Caetano Veloso |
| | goodnight, dear | Young K |
| | I Say a Little Prayer | Aretha Franklin |
| | Have You Ever Seen The Rain | Creedence Clearwater Revival |
| | Perfect Day | Lou Reed |
| | Wrecked | Imagine Dragons |
| | Merry Christmas Mr. Lawrence | Ryuichi Sakamoto |
| | Patience | Guns N' Roses |
| Power | Will Power | アトラスサウンドチーム |
| | Capabilities Unseen (feat. L) | Void_Chords |
| | GODS | League of Legends |
| | Big in Japan | Guano Apes |
| | You Oughta Know - 2015 Remaster | Alanis Morissette |
| | Fight Song | Rachel Platten |
| | The Upsetters | First Breath After Coma |
| | Killing In The Name | Rage Against The Machine |
| | Desire | MEG MYERS |
| | Toxicity | System Of A Down |
| | All I Want | The Offspring |
| | No One Knows | Queens of the Stone Age |
| | My Hero | Foo Fighters |
| | Deal With It (feat. Kelis) | Ashnikko |
| | Bring Me To Life | Evanescence |
| | Differently | Marian Hill |
| | The Rat | The Walkmen |
| | Under Pressure - Remastered 2011 | Queen |

| | | |
|---|---|---|
| | It's My Life | Bon Jovi |
| | Starlight | Muse |
| | Take Me Out | Franz Ferdinand |
| | Where U @ | Slow J |
| | Highway to Hell | AC/DC |
| | Sound of Madness | Shinedown |
| | Dia De Folga | Ana Moura |
| | Eu vi este povo a lutar (Confederação) | José Mário Branco |
| | Shout Out to My Ex | Little Mix |
| | Juice | Lizzo |
| | A Noite Dos Alquimistas | Fausto |
| | A Memória Dos Dias | Fausto |
| | Dog Days Are Over | Florence + The Machine |
| | Unstoppable | Sia |
| | Legions of Doom | Audiomachine |
| | Strength of a Thousand Men | Two Steps from Hell |
| | Symphony No. 9 in E Minor, Op. 95 "From the New World": IV. Allegro con fuoco | Antonín Dvořák |
| | No Cars Go | Arcade Fire |
| | Gila Monster | King Gizzard & The Lizard Wizard |
| | Personal Jesus | Depeche Mode |
| Sadness | John Wayne Gacy, Jr. | Sufjan Stevens |
| | Sol Negro | Maria Bethânia |
| | Que O Amor Te Salve Nesta Noite Escura - Ao Vivo | Pedro Abrunhosa |
| | Moonlight Sonata (1st Movement) | Rousseau |
| | Moonlight Sonata, Opus 27 No. 2, 1st Movement | Royal Symphony Orchestra |
| | Against the Wind | Various Artists |
| | Sorriso | Diogo Piçarra |
| | One More Light | Linkin Park |
| | Minha mãe | José Afonso |
| | True Love Waits | Radiohead |
| | Irene | Rodrigo Amarante |
| | Head Above Water | Avril Lavigne |
| | Ó gente da minha terra | Mariza |
| | Para Os Braços Da Minha Mãe | Pedro Abrunhosa |
| | A Gente Vai Continuar | Jorge Palma |

| | | |
|---|---|---|
| | On the Nature of Daylight | Max Richter |
| | Lag Fyrir Ömmu | Ólafur Arnalds |
| | Tears in Heaven | Eric Clapton |
| | Love You Goodbye | One Direction |
| | Two Cousins | Slow Club |
| | Will of the Scribes - Acoustic | Darren Korb |
| | Control | Zoe Wees |
| | Hurt | Johnny Cash |
| | Dumbledore's Farewell | Nicholas Hooper |
| | Sweet Child O Mine | Various Artists |
| | How to Save a Life | The Fray |
| | The Galaxy Last Moment!! A Phenomenally Awesome Guy | Dragon Ball |
| | Adagio in G Minor "Albinoni's Adagio" | Berliner Philharmoniker |
| | Ne me quitte pas | Jacques Brel |
| | sangue do meu sangue | Salvador Sobral |
| | DEATH | Melanie Martinez |
| | Used to Be | Weyes Blood |
| | Balada Da Despedida Do 5º Ano Jurídico 88/89 | Grupo de Fados Coimbra _ Toada Coimbrã |
| | To Build A Home | The Cinematic Orchestra |
| | She Wants | Metronomy |
| | Hope There's Someone | Antony and the Johnsons |
| | Visions of Gideon | Various Artists |
| | Young And Beautiful | Lana Del Rey |
| | Moving On | Kodaline |
| Tenderness | Guarda-me Esta Noite | Valter Lobo |
| | Hoppípolla - Live | Sigur Rós |
| | Better Together | Jack Johnson |
| | Hoppípolla | Sigur Rós |
| | Just Breathe | Pearl Jam |
| | Across the Stars (Love Theme from "Star Wars: Attack of the Clones") | John Williams |
| | Sun & Moon | Two Steps from Hell |
| | Winter Bear | V |
| | Kiss Me | Sixpence None The Richer |
| | Waste | Rhye |
| | Ocean of Tears | Caroline Polachek |

| | | |
|---|---|---|
| | A Fleeting Dream | SQUARE ENIX MUSIC |
| | Flamingo | Michel Petrucciani |
| | As | Becca Stevens |
| | I Want to Write You a Song | One Direction |
| | Só Tinha De Ser Com Você | Elis Regina |
| | What A Difference A Day Made | Jamie Cullum |
| | All Smiles | Jess Penner |
| | BB (Garupa de Moto Amarela) | Tim Bernardes |
| | Seaside | The Kooks |
| | Wind Of Change | Scorpions |
| | Ténpu Ki Bai | Mayra Andrade |
| | Canto Do Povo De Um Lugar - Remixed Original Album | Caetano Veloso |
| | Un Vestido Y Un Amor | Caetano Veloso |
| | Dead Hearts | Stars |
| | I Will Follow You into the Dark | Death Cab for Cutie |
| | Vagalumes | POLLO |
| | Careful (From The Original Motion Picture "Magic Mike's Last Dance") | Lucky Daye |
| | Toxic | Yael Naim |
| | World Spins Madly On | The Weepies |
| | Here I Go Again - 1987 Version; 2017 Remaster | Whitesnake |
| | Don't Watch Me Cry | Jorja Smith |
| | Jigsaw Falling Into Place | Radiohead |
| | Magic | Coldplay |
| | Gotten (feat. Adam Levine) | Slash |
| | Wonderful Tonight | Eric Clapton |
| | Valentine | Laufey |
| | Stay Ready (What A Life) | Jhené Aiko |
| | Love Of My Life - Remastered 2011 | Queen |
| | Ballerina | Yehezkel Raz |
| Tension | Angel | Sarah McLachlan |
| | The Imitation Game | Alexandre Desplat |
| | No Time To Die | Billie Eilish |
| | Canción Sin Miedo | Vivir Quintana |
| | Succession (Main Title Theme) - Extended Intro Version | Nicholas Britell |
| | Fuck Authority | Pennywise |

| | | |
|---|---|---|
| | Not Afraid | Eminem |
| | Intro: Never Mind | BTS |
| | Time - 2011 Remastered Version | Pink Floyd |
| | All Eyes On Me | Bo Burnham |
| | Time | Hans Zimmer |
| | Roman's Beat - "Hearts" | Nicholas Britell |
| | Ouvi Dizer | Ornatos Violeta |
| | The Poet's Death | Mazgani |
| | Everything In Its Right Place | Radiohead |
| | Another Brick In The Wall - 2001 Remastered Version | Pink Floyd |
| | Non, je ne regrette rien | Édith Piaf |
| | Numb | Linkin Park |
| | abcdefu | GAYLE |
| | Psyco - From the Movie Psyco | Blackround Philharmonic Orchestra |
| | The Phantom of the Opera - Single Album Version | Andrew Lloyd Webber |
| | MILK OF THE SIREN | Melanie Martinez |
| | Ligeti: Requiem: II. Kyrie. Molto espressivo | György Ligeti |
| | Hope There's Someone | Antony and the Johnsons |
| | Please Don't | Graveyard |
| | Hustle Bones | Death Grips |
| | Nem Deus nem senhor | José Mário Branco |
| | Bunsen Burner | Ben Salisbury |
| | The Turing Test | Ben Salisbury |
| | Coward | Hans Zimmer |
| | Gekko | Various Artists |
| | Como Um Sonho Acordado | Fausto |
| | Os Vampiros | José Afonso |
| | The End | The Doors |
| | Painkiller | Judas Priest |
| | Goodbye - Theme from Dark, a Netflix Original Series | Apparat |
| | Plans We Made | Son Lux |
| | 背後からの使者〜巨像との戦い〜 | SIE Sound Team |
| Transcendence | Bohemian Rhapsody | Queen |
| | Tchaikovsky: Swan Lake, Op. 20, Act 2: No. 10, Scene. Moderato | Pyotr Ilyich Tchaikovsky |

| | | |
|---|---|---|
| | The Great Gig In The Sky - 2011 Remastered Version | Pink Floyd |
| | Donde Vas | Brigada Victor Jara |
| | Hino Sport Lisboa Benfica Stadium | NN Chants Benfica. Francisco1904 |
| | Fairytale Tree | Luftschloss |
| | Zombie (feat. Valerie Broussard) | ILLENIUM |
| | Lower Your Eyelids To Die With the Sun | M83 |
| | Sirene - Zen Baboon Remix | Paul Schulleri |
| | Gymnopédie No. 1 | Various Artists |
| | Arrival of the Birds | The Cinematic Orchestra |
| | From The Mouth Of Gabriel | Sufjan Stevens |
| | New Divide | Linkin Park |
| | La Partida | Victor Jara |
| | The Arena | Lindsey Stirling |
| | Cornfield Chase | Hans Zimmer |
| | Piano Concerto No. 1 in B-Flat Minor, Op. 23: I. Allegro non troppo e molto maestoso – Allegro con spirito - Live at Philharmonie, Berlin | Evgeny Kissin |
| | Numb | Elderbrook |
| | Perfect Guidance (Brazil) | Tjak |
| | Kônô | Bako Dagnon |
| | Intro | The xx |
| | Nuvole Bianche | Ludovico Einaudi |
| | The Best Is Yet To Come | Various Artists |
| | Now We Are Free | The Lyndhurst Orchestra |
| | The Mission | Ennio Morricone |
| | Wild Flower (with youjeen) | RM |
| | Paradise | Coldplay |
| | Hallelujah | Jeff Buckley |
| | You & Me - Flume Remix | Disclosure |
| | God's Gonna Cut You Down | Johnny Cash |
| | Sunrise | Coldplay |
| | Serenade for Strings in C major, Op. 48: I. Pezzo in forma di sonatina: Andante non troppo - Allegro moderato | Moscow Soloists |
| | Also Frightened | Animal Collective |
| | Atmosphere - 2020 Digital Remaster | Joy Division |
| | Chevaliers De Sangreal - From The Da Vinci Code Original Motion Picture Soundtrack | Hans Zimmer |

| | | |
|---|---|---|
| | A Way of Life | Hans Zimmer |
| | Old Pine | Ben Howard |
| | Symphony No. 2 in C minor - "Resurrection" / 5th Movement: Pesante | Gustav Mahler |
| | Shatter Me | Lindsey Stirling |
| | Day One (Interstellar Theme) | Hans Zimmer |
| Wonder | Nicaragua | Klô Pelgag |
| | Cold Cold Cold | Cage The Elephant |
| | Aisling Song | Various Artists |
| | Canção De Engate - Acoustic | Tiago Bettencourt |
| | Accidentally In Love - From "Shrek 2" Soundtrack | Counting Crows |
| | プロローグ～古えの地へ～ | SIE Sound Team |
| | Healer | DAY6 |
| | I Can See Clearly Now | Johnny Nash |
| | La valse d'Amélie | Yann Tiersen |
| | A New Hope and End Credits | John Williams |
| | Theme From Jurassic Park | John Williams |
| | Mystery of Love | Sufjan Stevens |
| | Cello Suite No. 1 in G Major, BWV 1007: I. Prélude | Johann Sebastian Bach |
| | Downtown | Allie X |
| | Hurricane | Fleurie |
| | aBem | Rui Massena |
| | Experience | Ludovico Einaudi |
| | Screwed (feat. Zoë Kravitz) | Janelle Monáe |
| | Wanderlust - Ratatat Remix | Björk |
| | Don't Stop Me Now - Remastered 2011 | Queen |
| | Stand by Me | Playing For Change |
| | oh baby | LCD Soundsystem |
| | The Universal | Blur |
| | I'm Good (Blue) | David Guetta |
| | Blackbird - Remastered 2009 | The Beatles |
| | Gabriel's Oboe | Ennio Morricone |
| | Paper Rings | Taylor Swift |
| | Númenor | Bear McCreary |
| | Symphony No. 5 In E Minor, Op. 64, TH.29: II. Andante cantabile, con alcuna licenza - Moderato con anima | Pyotr Ilyich Tchaikovsky |

| | | |
|---|---|---|
| | A Próxima Viagem | Cassete Pirata |
| | Cinema Paradiso (Main Theme) | Ennio Morricone |
| | November Rain - 2022 Version | Guns N' Roses |
| | Comfortably Numb - 2011 Remastered Version | Pink Floyd |
| | Danza Kuduro | Various Artists |
| | If I Lose Myself - Alesso vs OneRepublic | Alesso |
| | Cantiga para quem sonha | Luiz Goes |
| | Walk Of Life | Dire Straits |
| | The Heart Asks Pleasure First | Michael Nyman |
| | Kiss Me | Sixpence None The Richer |

### Decoding music vs. noise - a sanity check

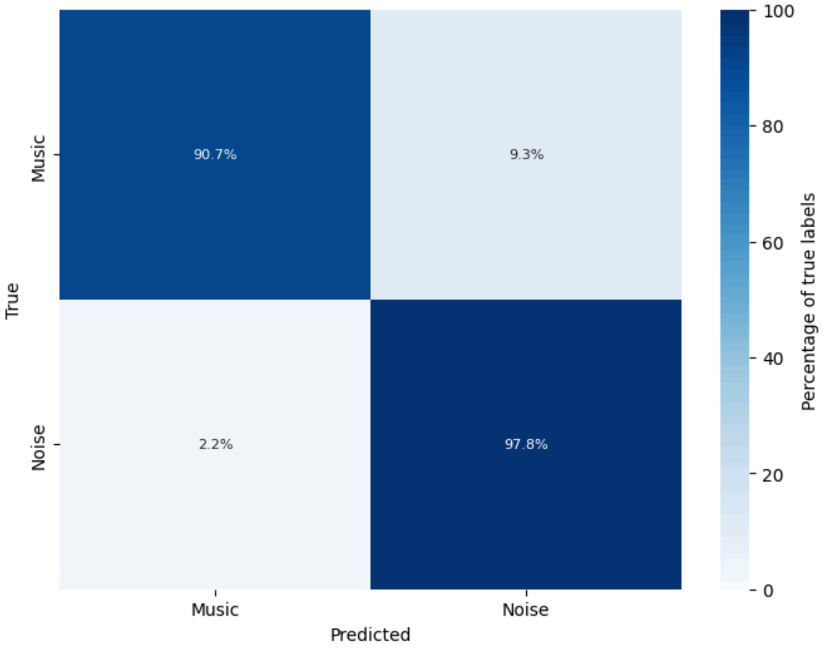


**Figure S2 - Group-level confusion matrix showing the results of the testing phase across the 20 participants while considering two classes: Music (including all emotions) and Noise. This analysis served as a sanity check for the features extracted. The overall balanced accuracy was 94.2% ± 3.2%, while the chance level sits at 50% in this case.**

### Decoding the three second-order emotion factors - sublimity, vitality, unease

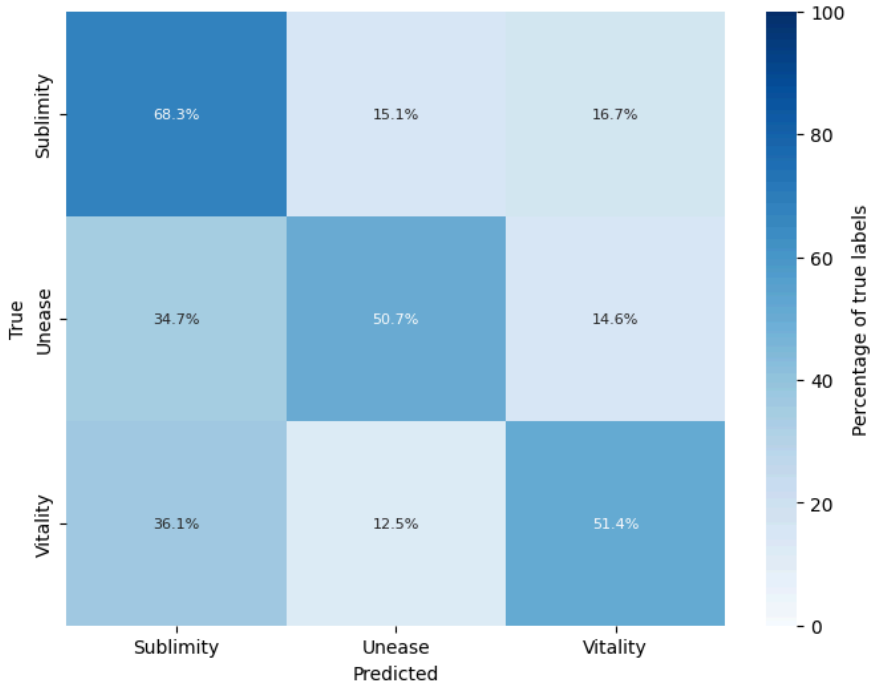


**Figure S3 - Group-level confusion matrix showing the results of the testing phase across the 20 participants while considering three classes, corresponding to the second-level factors of the GEMS model: Sublimity, Vitality, and Unease. The overall balanced accuracy was 56.8% ± 5.5%, while the chance level sits at 33.3% in this case.**

### Activation patterns per emotion in three core regions

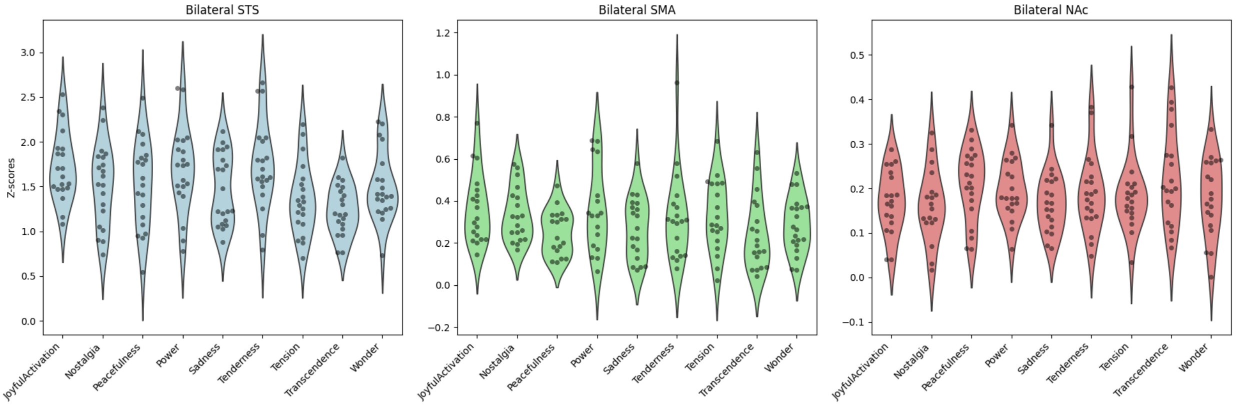


**Figure S4 - Distribution of activation values across the nine emotions in three core regions that responded significantly in the analysis reported in Figure 3: bilateral superior temporal sulcus (STS), supplementary motor area (SMA), and nucleus accumbens (NAc). No significant differences in activity were found across subjects between any emotions.**

### fMRIPrep methods

Results included in this manuscript come from preprocessing performed using fMRIPrep 23.1.2 (Esteban et al. (2019); Esteban et al. (2018); RRID:SCR_016216), which is based on Nipype 1.8.6 (K. Gorgolewski et al. (2011); K. J. Gorgolewski et al. (2018); RRID:SCR_002502).

#### Preprocessing of B0 inhomogeneity mappings

A total of 3 fieldmaps were found available within the input BIDS structure for this particular subject. A B0-nonuniformity map (or fieldmap) was estimated based on two (or more) echo-planar imaging (EPI) references with topup (Andersson, Skare, and Ashburner (2003); FSL None).

#### Anatomical data preprocessing

A total of 1 T1-weighted (T1w) images were found within the input BIDS dataset.The T1-weighted (T1w) image was corrected for intensity non-uniformity (INU) with N4BiasFieldCorrection (Tustison et al. 2010), distributed with ANTs (version unknown) (Avants et al. 2008, RRID:SCR_004757), and used as T1w-reference throughout the workflow. The T1w-reference was then skull-stripped with a Nipype implementation of the antsBrainExtraction.sh workflow (from ANTs), using OASIS30ANTs as target template. Brain tissue segmentation of cerebrospinal fluid (CSF), white-matter (WM) and gray-matter (GM) was performed on the brain-extracted T1w using fast (FSL (version unknown), RRID:SCR_002823, Zhang, Brady, and Smith 2001). Brain surfaces were reconstructed using recon-all (FreeSurfer 7.3.2, RRID:SCR_001847, Dale, Fischl, and Sereno 1999), and the brain mask estimated previously was refined with a custom variation of the method to reconcile ANTs-derived and FreeSurfer-derived segmentations of the cortical gray-matter of Mindboggle (RRID:SCR_002438, Klein et al. 2017). Volume-based spatial normalization to one standard space (MNI152NLin2009cAsym) was performed through nonlinear registration with antsRegistration (ANTs (version unknown)), using brain-extracted versions of both T1w reference and the T1w template. The following template was were selected for spatial normalization and accessed with TemplateFlow (23.0.0, Ciric et al. 2022): ICBM 152 Nonlinear Asymmetrical template version 2009c [Fonov et al. (2009), RRID:SCR_008796; TemplateFlow ID: MNI152NLin2009cAsym].

#### Functional data preprocessing

For each of the 4 BOLD runs found per subject (across all tasks and sessions), the following preprocessing was performed. First, a reference volume and its skull-stripped version were generated using a custom methodology of fMRIPrep. Head-motion parameters with respect to the BOLD reference (transformation matrices, and six corresponding rotation and translation parameters) are estimated before any spatiotemporal filtering using mcflirt (FSL , Jenkinson et al. 2002). The estimated fieldmap was then aligned with rigid-registration to the target EPI (echo-planar imaging) reference run. The field coefficients were mapped on to the reference EPI using the transform. BOLD runs were slice-time corrected to 0.446s (0.5 of slice acquisition range 0s-0.892s) using 3dTshift from AFNI (Cox and Hyde 1997, RRID:SCR_005927). The BOLD reference was then co-registered to the T1w reference using bbregister (FreeSurfer) which implements boundary-based registration (Greve and Fischl 2009). Co-registration was configured with six degrees of freedom. Several confounding time-series were calculated based on the preprocessed BOLD: framewise displacement (FD), DVARS and three region-wise global signals. FD was computed using two formulations following Power (absolute sum of relative motions, Power et al. (2014)) and Jenkinson (relative root mean square displacement between affines, Jenkinson et al. (2002)). FD and DVARS are calculated for each functional run, both using their implementations in Nipype (following the definitions by Power et al. 2014). The three global signals are extracted within the CSF, the WM, and the whole-brain masks. Additionally, a set of physiological regressors were extracted to allow for component-based noise correction (CompCor, Behzadi et al. 2007). Principal components are estimated after high-pass filtering the preprocessed BOLD time-series (using a discrete cosine filter with 128s cut-off) for the two CompCor variants: temporal (tCompCor) and anatomical (aCompCor). tCompCor components are then calculated from the top 2% variable voxels within the brain mask. For aCompCor, three probabilistic masks (CSF, WM and combined CSF+WM) are generated in anatomical space. The implementation differs from that of Behzadi et al. in that instead of eroding the masks by 2 pixels on BOLD space, a mask of pixels that likely contain a volume fraction of GM is subtracted from the aCompCor masks. This mask is obtained by dilating a GM mask extracted from the FreeSurfer's aseg segmentation, and it ensures components are not extracted from voxels containing a minimal fraction of GM. Finally, these masks are resampled into BOLD space and binarized by thresholding at 0.99 (as in the original implementation). Components are also calculated separately within the WM and CSF masks. For each CompCor decomposition, the k components with the largest singular values are retained, such that the retained components' time series are sufficient to explain 50 percent of variance across the nuisance mask (CSF, WM, combined, or temporal). The remaining components are dropped from consideration. The head-motion estimates calculated in the correction step were also placed within the corresponding confounds file. The confound time series derived from head motion estimates and global signals were expanded with the inclusion of temporal derivatives and quadratic terms for each (Satterthwaite et al. 2013). Frames that exceeded a threshold of 0.5 mm FD or 1.5 standardized DVARS were annotated as motion outliers. Additional nuisance timeseries are calculated by means of principal components analysis of the signal found within a thin band (crown) of voxels around the edge of the brain, as proposed by (Patriat, Reynolds, and Birn 2017). The BOLD time-series were resampled into standard space, generating a preprocessed BOLD run in MNI152NLin2009cAsym space. First, a reference volume and its skull-stripped version were generated using a custom methodology of fMRIPrep. All resamplings can be performed with a single interpolation step by composing all the pertinent transformations (i.e. head-motion transform matrices, susceptibility distortion correction when available, and co-registrations to anatomical and output spaces). Gridded (volumetric) resamplings were performed using antsApplyTransforms (ANTs), configured with Lanczos interpolation to minimize the smoothing effects of other kernels (Lanczos 1964). Non-gridded (surface) resamplings were performed using mri_vol2surf (FreeSurfer).

#### Functional data preprocessing

For each of the 4 BOLD runs found per subject (across all tasks and sessions), the following preprocessing was performed. First, a reference volume and its skull-stripped version were generated using a custom methodology of fMRIPrep. Head-motion parameters with respect to the BOLD reference (transformation matrices, and six corresponding rotation and translation parameters) are estimated before any spatiotemporal filtering using mcflirt (FSL , Jenkinson et al. 2002). The estimated fieldmap was then aligned with rigid-registration to the target EPI (echo-planar imaging) reference run. The field coefficients were mapped on to the reference EPI using the transform. BOLD runs were slice-time corrected to 0.445s (0.5 of slice acquisition range 0s-0.89s) using 3dTshift from AFNI (Cox and Hyde 1997, RRID:SCR_005927). The BOLD reference was then co-registered to the T1w reference using bbregister (FreeSurfer) which implements boundary-based registration (Greve and Fischl 2009). Co-registration was configured with six degrees of freedom. Several confounding time-series were calculated based on the preprocessed BOLD: framewise displacement (FD), DVARS and three region-wise global signals. FD was computed using two formulations following Power (absolute sum of relative motions, Power et al. (2014)) and Jenkinson (relative root mean square displacement between affines, Jenkinson et al. (2002)). FD and DVARS are calculated for each functional run, both using their implementations in Nipype (following the definitions by Power et al. 2014). The three global signals are extracted within the CSF, the WM, and the whole-brain masks.

Additionally, a set of physiological regressors were extracted to allow for component-based noise correction (CompCor, Behzadi et al. 2007). Principal components are estimated after high-pass filtering the preprocessed BOLD time-series (using a discrete cosine filter with 128s cut-off) for the two CompCor variants: temporal (tCompCor) and anatomical (aCompCor). tCompCor components are then calculated from the top 2% variable voxels within the brain mask. For aCompCor, three probabilistic masks (CSF, WM and combined CSF+WM) are generated in anatomical space. The implementation differs from that of Behzadi et al. in that instead of eroding the masks by 2 pixels on BOLD space, a mask of pixels that likely contain a volume fraction of GM is subtracted from the aCompCor masks. This mask is obtained by dilating a GM mask extracted from the FreeSurfer's aseg segmentation, and it ensures components are not extracted from voxels containing a minimal fraction of GM. Finally, these masks are resampled into BOLD space and binarized by thresholding at 0.99 (as in the original implementation). Components are also calculated separately within the WM and CSF masks. For each CompCor decomposition, the k components with the largest singular values are retained, such that the retained components' time series are sufficient to explain 50 percent of variance across the nuisance mask (CSF, WM, combined, or temporal). The remaining components are dropped from consideration. The head-motion estimates calculated in the correction step were also placed within the corresponding confounds file. The confound time series derived from head motion estimates and global signals were expanded with the inclusion of temporal derivatives and quadratic terms for each (Satterthwaite et al. 2013). Frames that exceeded a threshold of 0.5 mm FD or 1.5 standardized DVARS were annotated as motion outliers. Additional nuisance timeseries are calculated by means of principal components analysis of the signal found within a thin band (crown) of voxels around the edge of the brain, as proposed by (Patriat, Reynolds, and Birn 2017). The BOLD time-series were resampled into standard space, generating a preprocessed BOLD run in MNI152NLin2009cAsym space. First, a reference volume and its skull-stripped version were generated using a custom methodology of fMRIPrep. All resamplings can be performed with a single interpolation step by composing all the pertinent transformations (i.e. head-motion transform matrices, susceptibility distortion correction when available, and co-registrations to anatomical and output spaces). Gridded (volumetric) resamplings were performed using antsApplyTransforms (ANTs), configured with Lanczos interpolation to minimize the smoothing effects of other kernels (Lanczos 1964). Non-gridded (surface) resamplings were performed using mri_vol2surf (FreeSurfer).

Many internal operations of fMRIPrep use Nilearn 0.10.1 (Abraham et al. 2014, RRID:SCR_001362), mostly within the functional processing workflow. For more details of the pipeline, see the section corresponding to workflows in fMRIPrep's documentation.

#### Copyright Waiver

The above boilerplate text was automatically generated by fMRIPrep with the express intention that users should copy and paste this text into their manuscripts unchanged. It is released under the CC0 license.
